# Chronic AMPK inactivation slows SHH medulloblastoma progression by inhibiting mTORC1 signaling and depleting tumor stem cell populations

**DOI:** 10.1101/2021.12.09.471978

**Authors:** Taylor Dismuke, Daniel Shiloh Malawsky, Hedi Liu, Jay Brenman, Biplab Dasgupta, Andrey Tikunov, Timothy R. Gershon

**Author notes:** corresponding author, Corresponding author: Timothy Gershon, MD, PhD, Emory University School of Medicine, Emory Children’s Center room 256, Atlanta, GA 30322. contributed equally to the work (Taylor Dismuke, Daniel Shiloh Malawsky).

## Abstract

We show that inactivating AMPK *in vivo* in a genetic model of medulloblastoma depletes tumor stem cell populations and slows tumor progression. Medulloblastoma, the most common malignant pediatric brain tumor, grows as heterogenous communities comprising diverse types of tumor and stromal cells. Previously, we showed that different populations in medulloblastomas show different sensitivities to specific targeted therapies. To determine if specific populations depend on AMPK, we analyzed mice with AMPK-inactivated medulloblastomas. We engineered mice with conditional deletion of the AMPK catalytic subunits *Prkaa1* and *Prkaa2* and conditional expression *SmoM2*, an oncogenic *Smo* allele that hyperactivates Sonic Hedgehog (SHH) signaling. We compared these medulloblastomas to SmoM2-driven medulloblastomas in AMPK-intact mice. AMPK-inactivation slowed tumor growth and progression, allowing longer event-free survival (EFS). scRNA-seq showed that AMPK inactivation altered cellular heterogeneity, increasing differentiation, decreasing tumor stem cell populations and reducing glio-neuronal multipotency. Surprisingly, AMPK-inactivated tumors showed decreased mTORC1 activation and *Hk2* expression. Genetic *Hk2* deletion in SmoM2-medulloblastomas similarly decreased stem cell populations, implicating reduced aerobic glycolysis in the tumor-suppressive effect of AMPK inactivation. Our results show that AMPK inactivation impairs tumor growth through mechanisms that disproportionately affect tumor stem cell populations that have proved refractory to conventional therapies.

## Introduction

Medulloblastoma is the most common malignant pediatric brain tumor and recurrence after treatment is the major cause of morbidity for medulloblastoma patients. Medulloblastoma, is a heterogeneous group of cancers with 4 major subgroups [1–4], and individual tumors in each subgroup show cellular heterogeneity Cellular heterogeneity may contribute to recurrence by increasing overall robustness to overcome selective pressures of therapy, and by generating specific resistant populations that drive recurrence. Understanding how different conditions affect the diversity of cell types within medulloblastomas is critical to designing therapies that can reduce the chance of recurrent disease. We examined the how reducing metabolic adaptability alters cellular heterogeneity by disabling the intracellular energy sensor AMP-activated kinase (AMPK) in a genetic mouse model of SHH medulloblastoma and then analyzing tumor growth and cellular diversity.

SHH (Sonic Hedgehog) subgroup tumors, the most frequent subgroup of medulloblastoma, can be recapitulated in mice genetically engineered to hyperactivate SHH signaling in the brain, producing tumors that resemble human SHH medulloblastoma in site, pathology, gene expression and cellular heterogeneity [5–7]. Studies of SHH-driven medulloblastomas in mice show that cellular heterogeneity contributes to tumor recurrence as individual cell types within tumors show different sensitivity or resistance to specific therapies and cells that survive treatment drive recurrence [8, 9]. Tumor stem cells, marked by expression of OLIG2, are specifically resistant to cytotoxic treatments currently used in medulloblastoma therapy and initiate post-treatment recurrence [8]. Currently, 20% of patients with SHH subgroup medulloblastoma experience incurable recurrence despite optimal up-front therapy. Identifying ways to target therapy-resistant tumor stem cells may allow conventional treatment to be effective for more patients.

Tumor cells in medulloblastoma show a range of differentiation states that reflect the developmental progression in cerebellar development from undifferentiated stem cells to committed cerebellar granule neuron progenitors (CGNPs) to differentiated neurons(including unipolar brush cells (UBCs) and cerebellar granule neurons (CGNs) [10]. However, medulloblastomas also contain tumor-derived cells with glial differentiation, and this divergence from the neural lineage of CGNPs provides indirect evidence of medulloblastoma stem cells with glio-neuronal potency [9, 11].

AMPK is a multi-subunit complex that is a primary regulator of cellular energy homeostasis [12], and disrupting AMPK function may produce important anti-tumor effects [13]. AMPK-mediated coordination of energy metabolism may be particularly important in SHH medulloblastomas, because SHH induces specialized metabolic features that support malignant growth [14], including HK2-dependent aerobic glycolysis [15, 16], PPAR-γ -dependent lipogenesis [17], HIF1α stabilization [18] and mitochondrial fragmentation [19]. These metabolic adaptations may increase the need for AMPK to sense energy scarcity in order to maintain energy homeostasis.

While AMPK is activated by the tumor suppressor LKB1, many lines of evidence show that the role of AMPK in cancer is more complex [20]. In contrast to the multiple inactivating LKB1 mutations that have been found in diverse human cancers [21–23], mutations in AMPK subunits causing human cancer are unknown. Divergent roles of LKB1 and AMPK are demonstrated by the *Kras^G12D^* mouse lung tumor model, in which *Lkb1* deletion accelerates tumor growth while co-deletion of the two AMPK catalytic subunits *Prkaa1* and *Prkaa2* inhibits tumorigenesis [24]. Apart from operating downstream of LKB1, numerous studies show that AMPK functions to increases metabolic adaptability in normal cells within the brain [25, 26]. These functions that may also be advantageous in brain tumors [27, 28]. Importantly, AMPK can be inactivated in the mouse brain or whole body without producing discernable developmental abnormalities [28–30]. While chronic, sustained AMPK inactivation through *Prkab1/1* codeletion in astrocytes results in neurodegeneration in aged mice [26], therapies that transiently disrupt AMPK may be safe in pediatric patients with medulloblastoma [28, 31].

Prior studies show that AMPK acts in medulloblastoma to support tumor growth, as deletion of *Prkaa2*, the gene encoding the AMPK*α*2 catalytic domain slows tumor progression in mice engineered to develop SHH-driven medulloblastoma [31]. One process contributing to this effect is loss of AMPK-mediated phosphorylation of the zinc finger protein CNBP, which decreases the translation of Ornithine Decarboxylase (ODC) and the production of polyamines [31, 32]. However, the mechanisms through which polyamine production permits tumor growth are unresolved, and other AMPK-dependent mechanisms may also affect medulloblastoma growth. The repertoire of glycolytic genes expressed by medulloblastoma cells changes with differentiation state [14–16], and this variation suggest that specific types of tumor cells within medulloblastomas may depend on AMPK for different purposes.

To resolve AMPK function in different tumor cell subpopulations within medulloblastomas, we investigated the consequence of AMPK catalytic domain ablation through *Prkaa1 and Prkaa2* co-deletion in medulloblastomas driven by oncogenic, mutant *Smo*. We used scRNA-seq and immunohistochemistry to examine the impact of AMPK inactivation on different types of cells within medulloblastomas, and compared to *Hk2*-deleted medulloblastomas to probe the mechanistic role of altered metabolism. Our studies show that *Prkaa1/Prkaa2* co-deletion, like *Hk2* deletion, increased intra-tumoral differentiation. Notably, *Prkaa1/Prkaa2* co-deletion reduced the populations of OLIG2-expressing stem cells that drive recurrence after conventional therapy. These studies provide new mechanistic insight into how metabolic specialization supports the diverse communities of tumor cells that promote recurrence after therapy.

## Methods

### Mice

We crossed *SmoM2* mice (Jackson Labs, stock # 005130) with *GFAP-Cre* mice (Jackson Labs, stock # 004600), to generate *G-Smo* mice. We crossed *Prkaa1^loxP/loxP^/Prkaa2^loxP/loxP^* mice that harbor *loxP* sites flanking the coding regions of *Prkaa1 and Prkaa2* [29] with GFAP-Cre mice to generate *Gfap-Cre/Prkaa1^loxP/loxP^/Prkaa2^loxP/loxP^ (G-Cre^Ampk^)* mice. We then crossed *G-Cre^Ampk^* mice with *SmoM2* mice generate *G-Smo^Ampk^* mice with *Prkaa1/2-deleted* medulloblastomas. All mice were of species Mus musculus and crossed into the C57BL/6 background through at least five generations. All animal studies were carried out with the approval of the University of North Carolina Institutional Animal Care and Use Committee under protocols (19-098 and 21-011).

### Histology and immunohistochemistry

Mouse brains were processed, immunostained and quantitatively analyzed as previously described [9, 33, 34]. Primary antibodies used were: OLIG2 diluted 1:100 (Cell Marque, # 387R-14), SOX10 diluted 1:200 (Cell Signaling Technology, #7833S). Stained images were counterstained with DAPI, digitally imaged using an Aperio Scan Scope XT (Aperio) and subjected to automated cell counting using Tissue Studio (Definiens).

### Tissue Preparation for Drop-seq

Data from *G-Smo* and *G-Smo^Ampk^* tumors were obtained by Drop-seq in separate batches, as described below. The *G-Smo* data was previously published [35] and made publicly available, while the *G-Smo^Ampk^* data were newly obtained for this study. In both batches, mice were euthanized by decapitation under isoflurane anesthesia. The brain was divided along the sagittal midline and one half was processed for histology and a sample of tumor was dissected from the other half and processed for Drop-seq analysis. This sample was dissociated using the Papain Dissociation System (Worthington Biochemical) following the protocol used in previous studies ^[9, 36]^. Briefly, tumor samples were incubated in papain at 37 °C for 15 min, then triturated and the suspended cells were spun through a density gradient of ovomucoid inhibitor.

We resuspended pelleted cells in 1 mL HBSS with 6 g/L glucose and diluted in PBS-BSA solution to a concentration of 95-110 cells/μL. Barcoded Seq B Drop-seq beads (ChemGenes) were diluted in Drop-seq lysis buffer to a concentration between 95-110 beads/μL. Tumor cells were co-encapsulated with barcoded beads using FlowJEM brand PDMS devices as previously described [9]. All cells were processed within one hour of tissue dissociation. Droplet breakage and library preparation steps followed Drop-seq protocol V3.1 [37]. After PCR, amplified cDNA was subjected Ampure XP cleanup at 0.6x and 1x ratios to eliminate residual PCR primers and debris. found by the bioanalyzer electropherogram. 1. If PCR failed to generate adequate cDNA, the PCR was repeated with the 3rd round increased from 11 to 13 cycles.

For QC purposes, library pools consisting of the tagmented cDNA from 2,000 beads/run were prepared and sequenced to low depth (~2.5M reads/2K beads). We used the resulting data to assess library efficiency, including total read losses to PolyA regions, nonsense barcodes and adapter sequences as well as the quality and number of the transcriptomes captured. Passable runs contained 40-60% of reads associated with the top 80-100 barcodes found in 2,000 beads. Drop-seq runs passing QC were then prepared for high-depth sequencing on an Illumina Novaseq S2 flow cell.

### Processing of scRNA-seq data

Data analysis was performed using the Seurat R package version 3.1.1 [38]. Data were subjected to several filtering steps. First, only genes that were detected in at least 30 cells were considered, to prevent misaligned reads appearing as rare transcripts in the data. Cells were then filtered using specific QC criteria to limit the analysis to cells with transcriptomes that were well-characterized and not apoptotic.

We noted that *G-Smo* cells were sequenced at a greater depth than *G-Smo^Ampk^* cells which can introduce unwanted batch effects into the analysis. Consistent with best practices [39], we downsampled the *G-Smo* cells to 46.5% of their original depth so as to achieve similar sequencing depth between *G-Smo* and *G-Smo^Ampk^* cells prior to further filtering.

We filtered out putative cells with fewer than 500 detected RNA molecules (nCount) or 200 different genes (nFeature), as likely to represent ambient RNA. We filtered out putative cells with greater than 4 standard deviations above the median nCount or nFeature as likely to be doublets, improperly merged barcodes, or sequencing artifacts. We also filtered out putative cells with more than 10% mitochondrial transcripts which we suspected to be dying cells.

### Harmony analysis

To merge the previously published *G-Smo* control tumor data with the *G-Smo^Ampk^* tumor dataset, we used the Harmony algorithm [40]. First, data from both tumor types were analyzed in single SCTransform normalization and PCA steps. The Harmony algorithm then used the cells’ PCA coordinates and dataset identity to calculate new coordinates for each cell so as to minimize dataset dependence when applying clustering to the cells. This algorithm produced a dimensional reduction that was used as a PCA for the following steps of the analysis.

### scRNA-seq Data normalization, clustering, differential gene expression, and visualization

We normalized the data was normalized using the SCTransform as implemented in Seurat, then selected the top 3,000 most highly variable genes. We performed PCA on these highly variable genes using the RunPCA function. The number of PCs to be used in downstream analysis was chosen based on the elbow plot as implemented by Seurat. We then used the FindNeighbors and FindClusters functions to identify cell clusters in the data.

To identify differential genes between clusters of cells, we used Wilcoxon rank sum test to compare gene expression of cells within the cluster of interest to all cells outside that cluster as implemented by the FindMarkers function. Specific parameters for the genes to be analyzed based on their log fold change between the two compared groups and percent of cells expressing the gene in at least one of the groups are available in the data analysis code. Uniform Manifold Approximation and Projection was used to reduce the PCs to two dimensions for data visualization using the RunUMAP function.

### Cell-type identification

Following PCA and UMAP, we analyzed cluster-specific differential gene expression. Marker genes were identified based on cluster-specific differential gene expression. For this purpose, for each gene we calculated the fraction of cells within the cluster that expressed the gene (referred to in Supplementary Data 1 as pct1) and the fraction of cells outside the cluster that expressed the gene (referred to in Supplementary Data 1 as pct2), and ranked genes by the ratio of pct1:pct2. Genes with high pct1:pct2 ratio and high pct1 values were selected as leading cluster-specific markers. The expression patterns of these genes were then examined in publicly available scRNA-seq data describing gene expression during mouse development [41] and in our prior published mouse medulloblastoma scRNA-seq studies [9, 35, 42], to identify cell types.

## Results

### AMPK signaling in SHH medulloblastoma

To investigate a potential dependence of medulloblastoma on AMPK, we analyzed publicly available data from medulloblastomas resected from patients. We compared expression of *PRKAA1* in tumors from each of the 4 medulloblastoma subgroups using the R2: Genomics Analysis and Visualization Platform (http://r2.amc.nl). While expression was variable in each subgroup, SHH subgroup medulloblastomas showed significantly higher *PRKAA1* expression compared to each of the other tumor types (Fig. 1A), suggesting a role for AMPK activity in the growth of SHH medulloblastomas.

**Figure 1.**
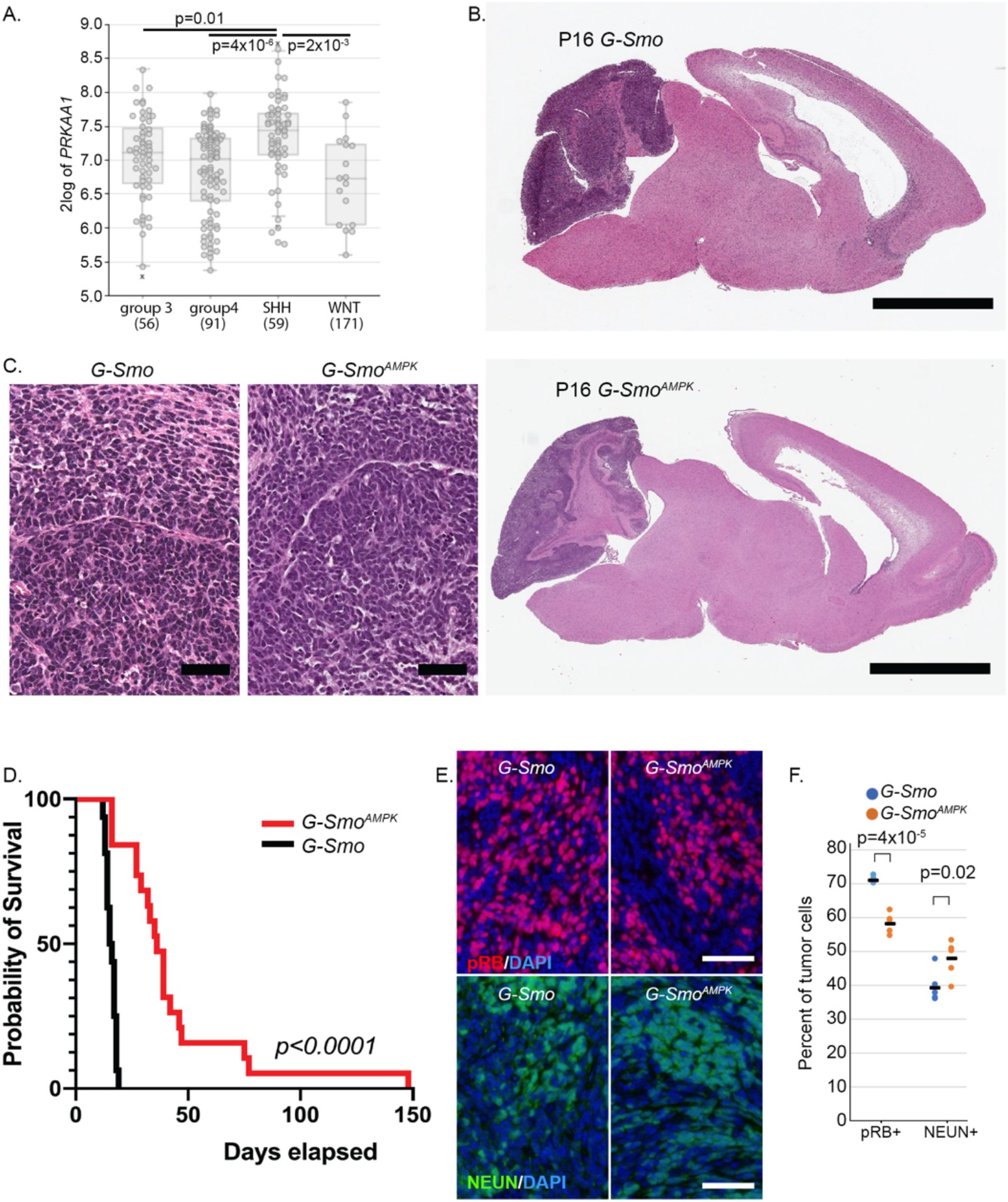
AMPK inactivation slows the progression of SHH-driven medulloblastoma. (A) *PRKAA1* mRNA expression in clinical medulloblastoma samples of indicated subgroups. (B,C) Representative sagittal brain sections from *G-Smo* and *G-Smo^Ampk^* mice show similar site of tumor formation and hypercellular tumor pathology. (D) Increased survival time in *G-Smo^Ampk^* mice compared to *G-Smo* controls with intact AMPK. Bars = 3 mm in (B) and 50 μm in (C). Survival times in (D) were compared by Log Rank test. (E) Representative IHe on sagittal sections of *G-Smo* and *G-Smo^Ampk^* medulloblastomas, showing pRB and NEUN expression. (F) quantification of the fractions of tumor cells expressing the indicated marker in replicate tumors of each genotype.

### AMPK inactivation in medulloblastomas increase survival of tumor-bearing mice

To test the importance of AMPK function in SHH medulloblastoma experimentally, we generated medulloblastoma-prone mice with CNS-specific *Prkaa1/Prkaa2* co-deletion. We first crossed *Gfap-Cre* mice that express Cre recombinase in CNS stem cells during development with *Prkaa1^loxP/loxP^/Prkaa2^loxP/loxP^* mice [29]. The resulting *G-Cre^Ampk^*mice were born at Mendelian ratios and showed showed normal survival and fertility, with no overt behavior changes, consistent with our prior studies [29]. Next, we bred *Prkaa1^loxP/loxP^/Prkaa2^loxP/loxP^* mice with the *SmoM2* mouse line [43] to generate mice with Cre-conditional AMPK inactivation and Cre-conditional SHH-pathway hyperactivation *(SmoM2^Ampk^)*. We then crossed *G-Cre^Ampk^* and *SmoM2^Ampk^* lines to produce *G-Smo^Ampk^* mice that featured AMPK inactivation, and SHH hyperactivation throughout the brain.

Brain-wide SHH hyperactivation in *G-Smo* mice produces medulloblastoma with 100% penetrance by postnatal day 10 (P10) and no other brain tumors [44]. Similar to *G-Smo* mice, all *G-Smo^Ampk^* mice developed medulloblastoma (Fig. 1B,C). In the absence of treatment, all *G-Smo* mice die of tumor progression by P22 [15, 45]. In contrast, untreated *G-Smo^Ampk^* mice showed significantly longer survival times, with median survival of 44 days in *G-Smo^Ampk^* compared to 16 days in controls (p<0.0001; Fig. 1D). These data show that AMPK inactivation in the *G-Smo* model does not prevent medulloblastoma formation, but impairs medulloblastoma progression, consistent with prior studies of isolated *Prkaa2* deletion in SHH medulloblastomas driven by the *SmoA1* allele.

Analysis of tumors from replicate *G-Smo^Ampk^* mice and *G-Smo* controls showed that AMPK inactivation altered the cellular composition of medulloblastomas, reducing proliferative populations and increasing differentiating populations. In medulloblastomas from both genotypes harvested at P15, we compared the fractions of cells expressing the proliferation marker phosphorylated RB (pRB) the differentiation marker NEUN, and the apoptosis marker cleaved Capase-3 (cC3; Fig.1E, Supplementary Fig. 1A). *G-Smo^Ampk^* medulloblastomas showed significantly lower proliferation index, (defined as pRB+ cells/total cells) and significantly higher differentiation index (defined as NEUN+ cells/total cells) cells (Fig. 1F). In contrast, the apoptotic rate trended higher in the *G-Smo^Ampk^* tumors with increased variability compared to controls, but was not significantly different (Supplementary Fig. 1B). These findings implicate decreased proliferation and increased differentiation rather than increased apoptosis in the slower progression of AMPK-inactivated tumors.

### AMPK inactivation alters medulloblastoma cellular heterogeneity and induces differentiation

To resolve changes in cellular heterogeneity within the AMPK-inactivated and control tumors, we used scRNA-seq to compare tumors from 5 replicate P15 *G-Smo^Ampk^* mice to 5 control medulloblastomas from replicate P15 *G-Smo* controls. We used the Harmony algorithm to integrate the scRNA-seq data from *G-Smo^Ampk^* and *G-Smo* tumors in a single analysis, generating a 2-dimensional UMAP projection in which individual cells were clustered with cells of similar gene expression. As in prior studies, this analysis defined both discrete clusters and a group of clusters with shared borders (Fig 2A). We determined the biological relevance of the clustering by generating cluster-specific differential gene expression profiles, comparing the expression by cells within the cluster to the expression by all cells outside the cluster (Supplementary Data 1). These gene expression profiles identified the cell type of each cluster.

**Figure 2.**
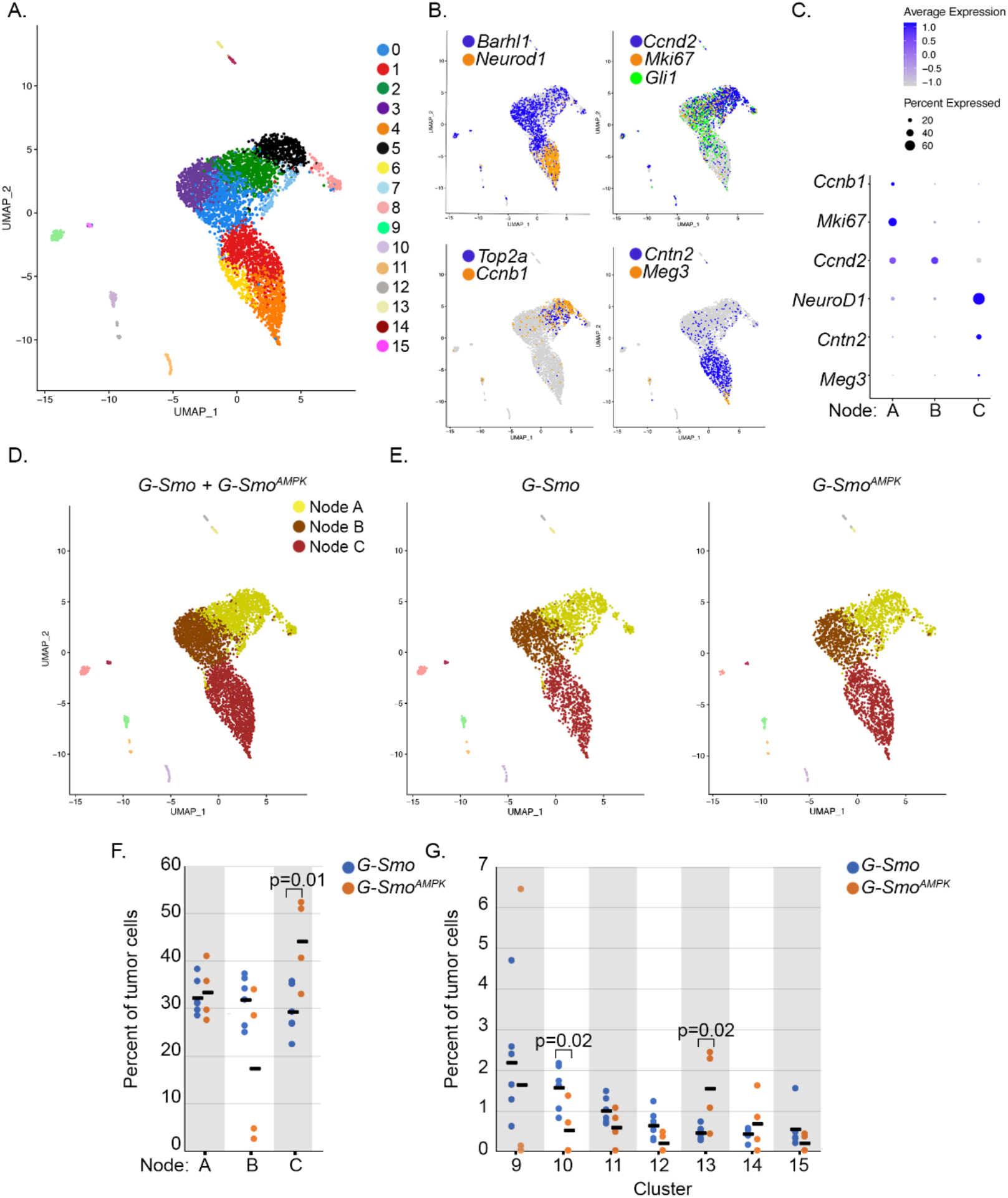
AMPK inactivation alters cellular heterogeneity in SHH driven medulloblastomas. (A) UMAP plot of all cells from *G-Smo* and *G-Smo^AMPK^* tumors, color-coded by cluster. Cells are localized according to their proximity in PCA space. (B) Feature plots showing expression of indicated proliferation or differentiation markers, color coded over the UMAP shown in (A). (C) Dot plot of indicated markers that division of tumor cells into Nodes A-C separates them by proliferation and differentiation states. (D) Nodes A-C projected onto the UMAP from (A). (E) UMAP plots deconvoluted by genotype, with Nodes A-C color coded. (F,G) Comparison of populations of (F) Nodes AC or (G) each stromal cluster in *G-Smo* and *G-Smo^Ampk^* tumors. Dots represent values for individual replicates and horizontal bars indicate the means. Student’s t-test was used to make pairwise comparison.

As in our prior studies, these methods identified the discrete clusters as different types of stromal cells typical of brain tissue including: astrocytes, oligodendrocytes, ependymal cells, myeloid cells, endothelial cells and fibroblasts (Table 1; Fig. 2A). Expression of CGNP markers *Barhl1* and *NeuroD1* and SHH transcription factor *Gli1* identified the 8 clusters in the multi-cluster complex as SHH-medulloblastoma cells (Fig. 2B). Cluster-specific gene expression analysis showed these clusters to represent a range of states that paralleled CGNP development, from proliferative, undifferentiated cells to non-proliferative cells at different stages of neural differentiation as described below and in Table 1. The full set of gene expression data with single cell resolution from *G-Smo^Ampk^* and *G-Smo* control tumors can be viewed at: https://malawsky.shinyapps.io/Gerhson_AMPK_analysis.

**Table 1.**
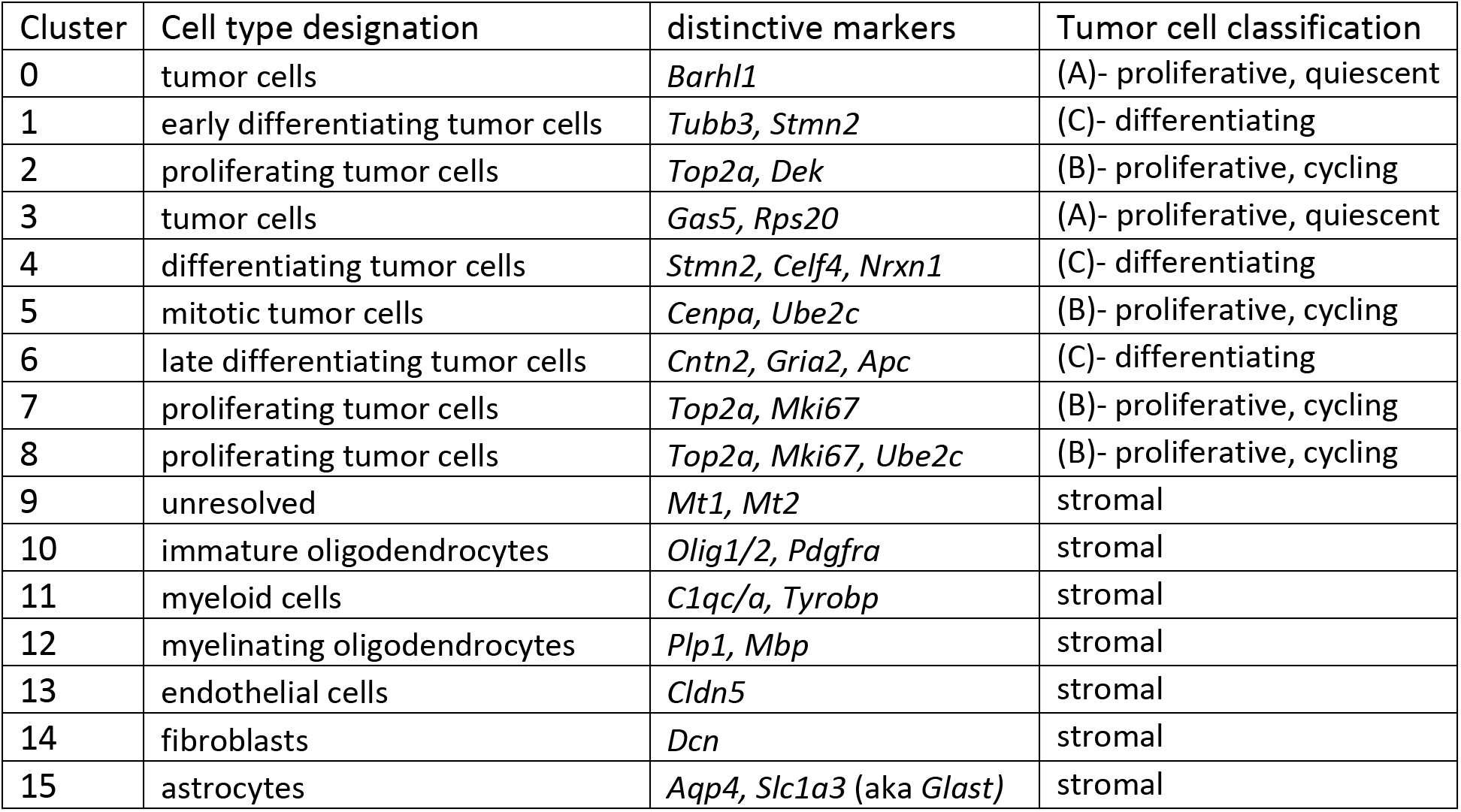
Identification of clusters as specific types of tumor and stromal cells.

To parse the multi-cluster complex of tumor cells, we designated tumor cell clusters as proliferative based by expression of proliferation markers *Mki67* and *Cyclin* genes and SHH target *Gli1* (Fig. 2B). We further characterized proliferative clusters as quiescent, cycling or M-phase enriched based on expression of S/G2/M marker *Top2a* [46] and mitotic marker *Ccnb1* [47] (Fig. 2B). The non-proliferative clusters showed successive expression of early to late differentiation markers *NeuroD1, Cntn2*, and *Grin2b* (Table 1; Fig 2B) with terminally differentiated neurons as the most differentiated cell type within this group [9, 45]. Using these markers, we classified the tumor cell clusters as (A) proliferative but quiescent, (B) proliferative and cycling, or (C) differentiating (Fig 2C,D; Table 1).

We disaggregated the cells by genotype (Fig. 2E) to compare the populations and gene expression patterns of type of cell in *G-Smo^Ampk^* and *G-Smo* control tumors. We analyzed the distribution of tumor cells across the differentiation spectrum by determining the number of cells from each replicate animal in Set A, B or C, normalized to the total number of cells from that animal (Fig. 2F). The distribution of tumor cells across the three sets was significantly different in *G-Smo^Ampk^* tumors (p=0.034; ANOVA), with differentiated cells (Set C) significantly increased (p=0.01; Student’s t-test), consistent with increased NEUN+ cells in *G-Smo^Ampk^* tumors detected by IHC.

Similarly, we compared the populations of each stromal cluster in *G-Smo^Ampk^* and *G-Smo* control tumors (Fig. 3G). We did not detect significant differences in most stromal populations, including astrocytes and myelinating oligodendrocytes which were both within the GFAP lineage and thus subject to *Prkaa1/2* deletion. However, immature oligodendrocytes (Cluster 10), which were within the GFAP lineage and thus also subject to *Prkaa1/2* deletion, were 3-fold less numerous in *G-Smo^Ampk^* tumors (p=0.02; Student’s t-test). Additionally, *G-Smo^Ampk^* tumors showed a 3-fold increase in endothelial cells (p=0.02; Student’s t-test), which are not within the GFAP lineage and therefore were not *Prkaa1/2-deleted*. AMPK inactivation thus increased differentiation in both the tumor and oligodendrocyte lineages and also produced a non-cell autonomous increase in endothelial cells.

**Figure 3.**
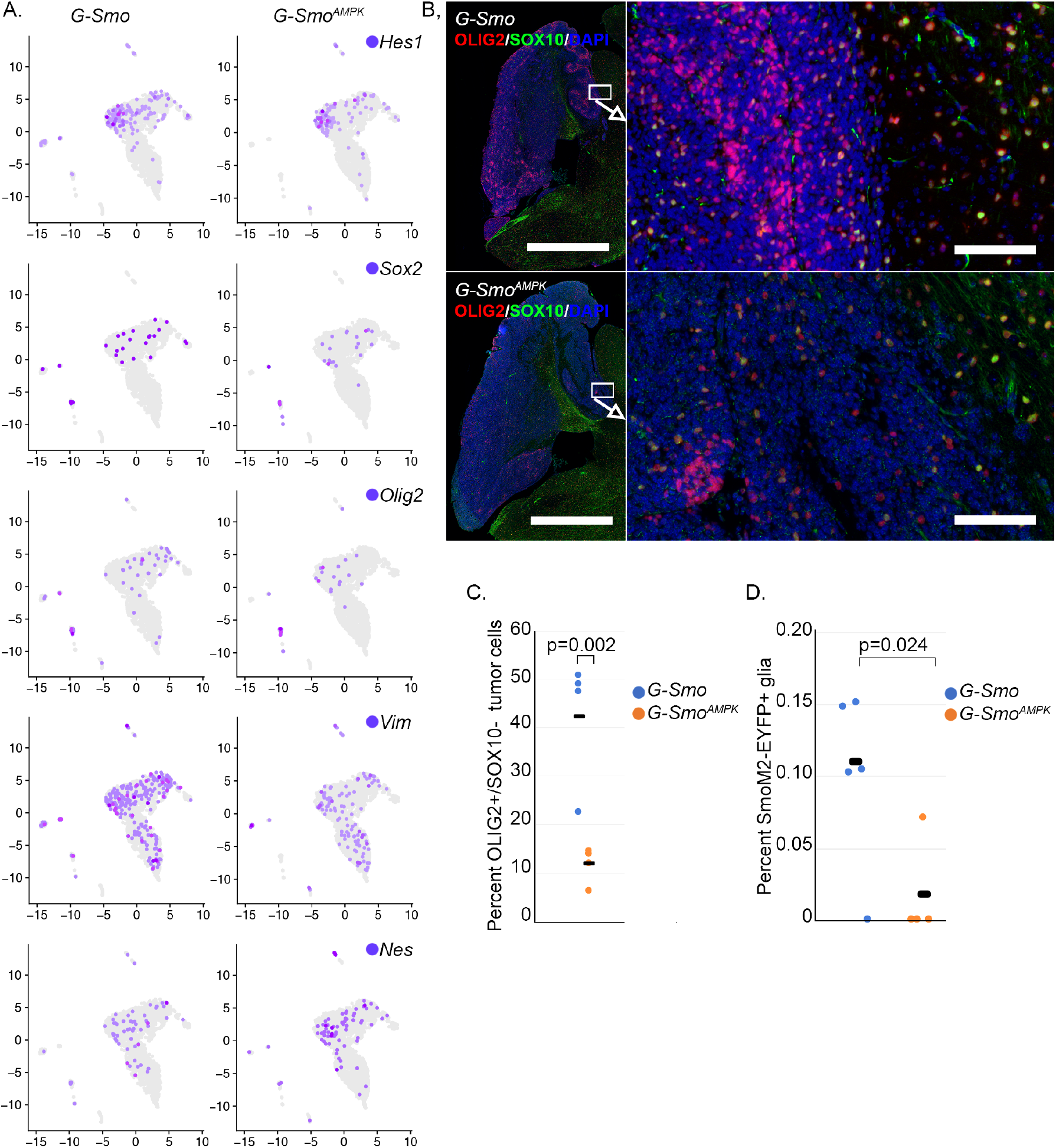
Decreased stem cell populations in *G-Smo^AMPK^* tumors. (A) Expression of indicated stem cell markers in the indicated genotypes, projected onto the disaggregated UMAPs. *Hes1, Sox2, Olig2* and *Vim* were significantly lower in *G-Smo^AMPK^* tumors, while Nes was not significantly different. (B) Representative OLIG2/SOX10 IHC in sagittal sections of tumors of indicated genotype. (C) Quantification of OLIG2/SOX10 IHC as in (B) in replicate samples of each genotype. (D) Quantification of SmoM2-YFP+ glial cells in the indicated genotypes. In (C) and (D), p-value was determined student’s t-test.

### AMPK-inactivated medulloblastomas show reduced OLIG2+ stem cells

Considering increased differentiation in AMPK-inactivated medulloblastomas and extended survival time, we determined if tumor stem cell populations, which remain undifferentiated and play an essential role in medulloblastoma progression, were reduced. We compared the populations of cells that expressed the stem cell markers markers *Hes1, Olig2, Sox2, Nes* and *Vim* in *G-Smo* versus *G-Smo^Ampk^* tumors (Fig. 3A). In the tumor cell clusters of Nodes A-C, *Olig2+* and *Vim+* cells were significantly reduced in *G-Smo^Ampk^* medulloblastomas, while *Sox2*+ cells and *Hes1+* cells showed trends toward reduced numbers that were not statistically significant.

To quantify stem cell populations using an alternative detection method, we analyzed OLIG2 protein expression, detected by IHC. As both tumor stem cells and oligodendrocytes express OLIG2, we used the oligodendrocyte marker SOX10 to distinguish between these two cells types, considering OLIG2+/SOX10-cells in medulloblastomas as tumor stem cells. These OLIG2+/SOX10-stem cells were increased in perivascular regions (Fig. 3B), consistent with prior descriptions of medulloblastoma stem cells [33, 48]. The fractions of OLIG2+/SOX10-cells were markedly reduced in *G-Smo^Ampk^* tumors, consistent with the differences in *Olig2+* populations in Nodes A-C in the scRNA-seq data (Fig. 3C). These scRNA-seq and immunohistochemistry studies consistently demonstrate decreased stem-like populations in AMPK-inactivated medulloblastomas.

### AMPK-inactivation disrupts tumor to glia trans-differentiation

Lineage tracing in mouse models of SHH medulloblastoma show that tumor cells trans-differentiate to generate tumor-derived glial populations [9, 11], and that these malignant glia play a supportive role in tumor growth [11]. Glial trans-differentiation requires stem-like pluripotency, and in light of the reduced stem-like population in *G-Smo^Ampk^* tumors, we compared glial trans-differentiation in *G-Smo^Ampk^* and *G-Smo* control tumors.

We traced lineage in the scRNA-seq data using the 3’*Yfp* sequence in the Cre-conditional *SmoM2* transgene. We previously used this method to show that subsets of astrocytes and oligodendrocytes derive from the tumor lineage in medulloblastomas that form in *Math1-Cre/SmoM2* mice [9]. In the *G-Smo* control tumors, we found *SmoM2-Yfp* expression in glial cells (Fig. 3D), which was expected since *Gfap-Cre* activates *SmoM2* in glio-neuronal stem cells of the developing brain that generate both glial and neuronal progeny. However, not all glia were *SmoM2-Yfp+*, indicating that *SmoM2* activation was not uniform in these populations, and this variation allowed us to quantify the glial cells of tumor lineage. *G-Smo^Ampk^* tumors showed significantly smaller *SmoM2-Yfp+* glial populations, indicating fewer tumor-derived glia (Fig. 3D, p-value 0.024). The decreased glial trans-differentiation provides functional evidence that stem cells were reduced in *G-Smo^Ampk^* tumors.

### Decreased mTORC1 activation in AMPK-inactivated medulloblastomas

To identify transcriptomic changes caused by AMPK inactivation, we compared gene expression patterns of *G-Smo^Ampk^* and *G-Smo* medulloblastoma cells within each cluster. We found multiple clusters showed consistent patterns of differential gene expression. We considered genes to be significantly down-regulated in *G-Smo^Ampk^* tumors if they met criteria of corrected p<0.05 and were expressed in fewer cells within the same cluster in *G-Smo^Ampk^* tumors compared to *G-Smo* tumors. With these criteria, we found 36 different genes significantly down-regulated in any of the 9 tumor cell clusters in *G-Smo^Ampk^* tumors. 14 of these 36 genes were down-regulated in at least 5 of 9 clusters, and 11 were down-regulated in at least 7 of 9 clusters (Fig. 4A,B; p<10^-15^ by extended hypergeometric test [49]). Similarly, 78 genes were up-regulated in any of the 9 tumor cell clusters in *G-Smo^Ampk^* tumors with criteria of corrected p<0.05 and expressed in more cells within the same cluster in *G-Smo^Ampk^* tumors compared to *G-Smo* tumors. 14 of these 78 genes were up-regulated in 5 of 9 clusters, and 8 were up-regulated in at least 7 of 9 clusters (p<0.001 by extended hypergeometric test). AMPK inactivation thus produced similar changes in gene expression across the heterogeneity of the tumors.

**Figure 4.**
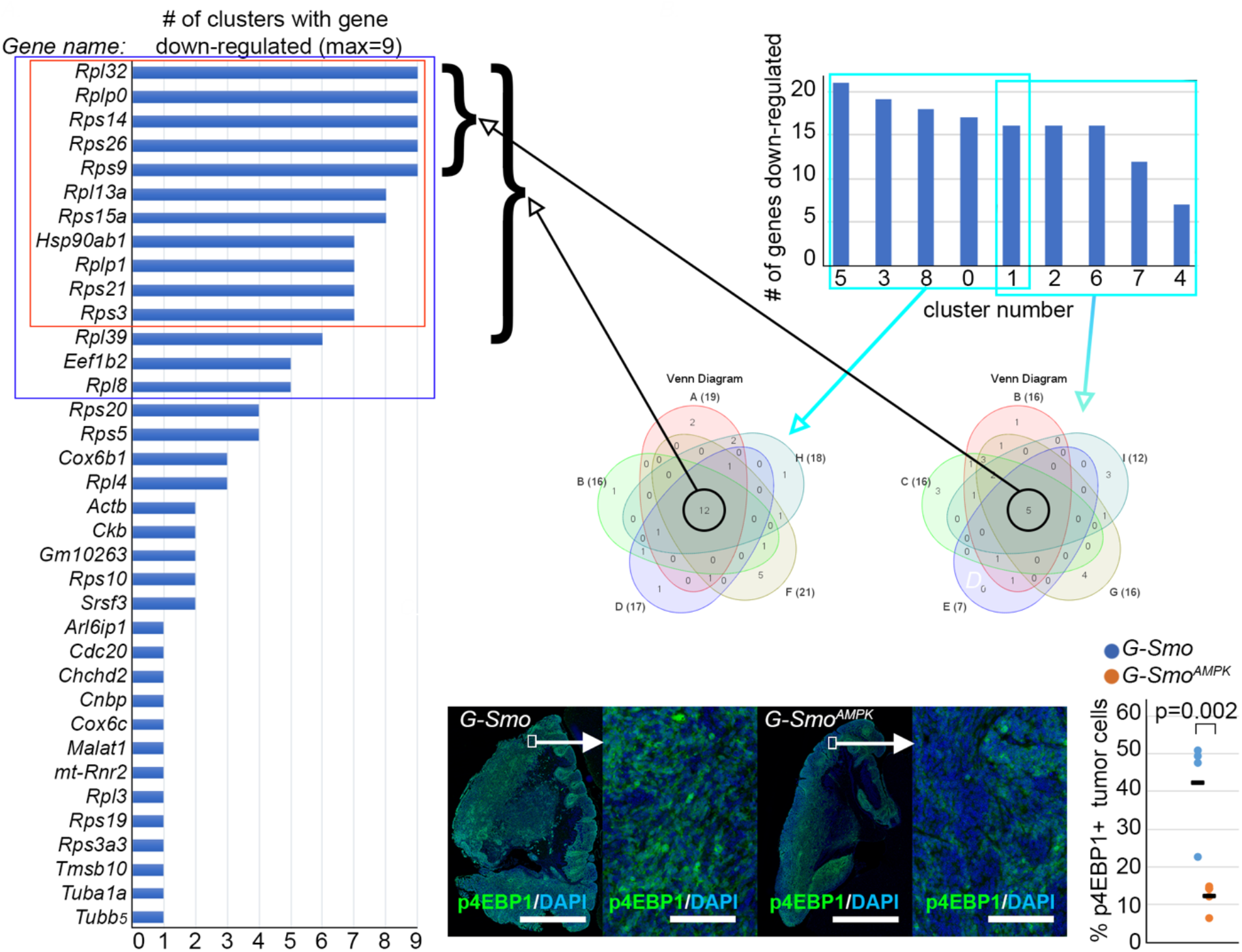
Decreased mTORC1 activity in *G-Smo^Ampk^* tumors. (A) Genes that are down-regulated in 1 or more clusters, ranked by number of clusters with differential expression. Red box marks the 11 genes down-regulated in 7/9 clusters. Blue box marks the 14 genes down regulated in 5/9 clusters. (B) Venn diagrams of the genes down-regulated in the 5/9 clusters with the highest number of differentially expressed genes, or in the 5/9 clusters with the fewest differentially expressed genes. Cluster 1 is included in both Venn diagrams as a point of consistency, as depicted in the graph. These Venn diagrams depict sets of 12 or 5 genes down-regulated in all included clusters and brackets show the indicated sets of genes commonly down-regulated genes in the list from (A). (C) Representative p4EBP1 IHC in sagittal sections of tumors of indicated genotype. (D) Quantification of p4EBP1 IHC as in (C) in replicate samples of each genotype.

GO analyses of the set of 14 genes down-regulated in 7/9 clusters in *G-Smo^Ampk^* tumors identified translation as the primary process statistically implicated (corrected p<4×10^-23^), driven by multiple ribosomal proteins and the translation elongation factor EEF1B2 (Fig. 4B). In contrast, GO analysis of the 14 genes up-regulated in 7/9 clusters in *G-Smo^Ampk^* tumors did not identify a statistically implicated process. We have previously found that down-regulation of translation-related genes in medulloblastomas correlates with decreased mTORC1 signaling [42], and we therefore analyzed whether mTORC1 signaling was altered in *G-Smo^Ampk^* tumors.

We assessed mTORC1 activation state by analyzing phosphorylation of the mTORC1 substrate 4EBP1 (p4EBP1), using immunohistochemistry. We quantified p4EBP1+ cells in tissue sections of *G-Smo^Ampk^* and control tumors (Fig. 4C). *G-Smo^Ampk^* tumors showed significantly fewer p4EBP1+ cells, and an F-score that integrated both the fraction of positive cells and the staining intensity was similarly decreased (Fig. 4D). Together the patterns of gene expression and p4EBP1 studies show reduced mTORC1 activation in *G-Smo^Ampk^* tumors compared to *G-Smo* control tumors. AMPK is known to inhibit mTORC1 [50–52], and the reduced mTORC1 activity in *G-Smo^Ampk^* tumors is opposite of the predicted effect of acute AMPK inhibition. This unexpected suppression of mTORC1 activity may result from a homeostatic response to the sustained AMPK inactivation in *G-Smo^Ampk^* tumors.

### Altered glycolytic gene expression in AMPK-inactivated medulloblastomas

We previously found that HK2-dependent aerobic glycolysis maintains medulloblastoma undifferentiated populations within SHH-driven medulloblastomas. In these prior studies, we crossed *G-Smo* mice with mice that harbor conditional *Hk2* deletion *(Hk2^loxP/loxP^)* to produce the genotype *Gfap-Cre/SmoM2/Hk2^loxP/loxP^ (G-Smo^Hk2^)*and found that the medulloblastoma that form in *G-Smo^Hk2^* mice showed increased differentiation, with regions of appropriately formed internal granule layer (IGL), increased event-free survival, and increased AMPK activation [15]. The correlation between Hk2 deletion, AMPK activation and increased in differentiation suggested that AMPK activation might be caused by Hk2 deletion and might induce differentiation. Alternatively, AMPK activation and increased differentiation might not be causally related. In support of this possibility, our current finding of increased differentiation in *G-Smo^Ampk^* tumors, however, demonstrates that differentiation does not require AMPK activation.

To test directly whether the differentiating effect of *Hk2* deletion in medulloblastoma requires AMPK function, we bred mice with medulloblastomas with both *Hk2* deletion and AMPK inactivation, genotype *Gfap-Cre/Prkaa1^loxP/loxP^/Prkaa2^loxP/loxP^/Hk2 ^loxP/loxP^ (G-Smo^Ampk/Hk2^)*. These *G-Smo^Ampk/Hk2^* mice developed medulloblastomas with 100% frequency, and the tumors resembled the previously described *G-Smo^Hk2^* medulloblastomas, marked by regions of differentiation (Fig. 5A) [15]. These co-deletion studies show that the increased differentiation phenotype caused by *Hk2* deletion does not require AMPK activity, and therefore that AMPK does not operate downstream of HK2-dependent glycolysis to regulate differentiation. In light of these findings, we examined the alternative possibility suggested by AMPK activation in *Hk2*-deleted tumors, that AMPK may operate upstream to induce HK2-dependent aerobic glycolysis.

**Figure 5.**
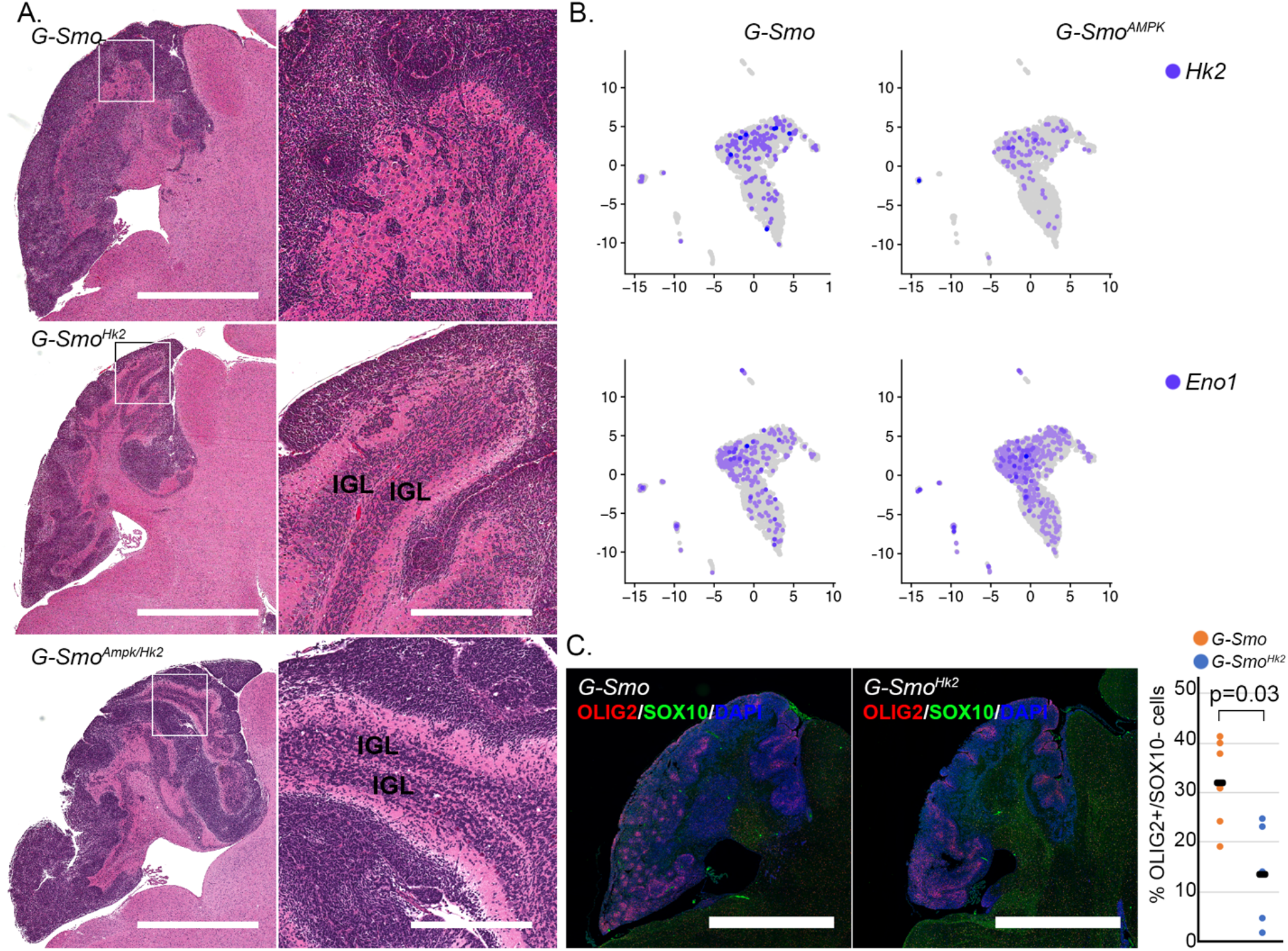
Defining the interaction of AMPK and HK2. (A) Representative H&E sections compare *G-Smo* control tumors to tumors with deletion of *Hk2* or co-deletion of *Hk2*,*Prkaa1* and *Prkaa2*. Increased differentiation in Hk2-deleted tumors is not rescued by co-deletion of *Prkaa1* and *Prkaa2*. IGL regions in *G-Smo^Hk2^* and *G-Smo^Ampk^* tumors are indicated. (B) scRNA-seq data showing decreased Hk2 and increased Eno1 in *M-Smo^Ampk^* tumors. (C) Representative OLIG2 immunofluorescence, comparing *G-Smo* and Hk2-deleted medulloblastomas, with quantification of replicate samples. P value determined by Student’s t-test.

We compared the number of cells expressing each gene in the glycolytic pathway in *G-Smo^Ampk^* medulloblastomas compared to *G-Smo* control tumors and found that *Hk2* and *Eno1* were differentially expressed in AMPK-deleted tumors. *Hk2+* cells were significantly reduced in *G-Smo^Ampk^* tumor cells (Fig. 5B), consistent with the hypothesis that AMPK activity modulates HK2 expression in SHH-driven medulloblastoma, as seen in prior studies of normal in muscle cells [53]. In contrast, cells expressing *Eno1*, which acts at the end of glycolysis, were increased in *G-Smo^Ampk^* tumors (Fig. 5B). AMPK inactivation thus induced specific changes in glycolytic gene expression, predicted to reduce entry of glucose into glycolysis and to increase the processing of glycolytic intermediates toward pyruvate.

### Similar stem cell changes in AMPK-inactivated and *Hk2-deleted* medulloblastomas

To determine if alterations in glycolysis may produce specific features of the AMPK-inactivation phenotype, we analyzed OLIG2+ populations in *Hk2*-deleted medulloblastomas. Comparing *G-Smo/Hk2^floxed^* mice to *G-Smo* controls we found that *Hk2*-deletion was sufficient to reduce the fraction of OLIG2+ tumor cells (Fig. 5C). Reduced tumor stem cell populations was thus a common feature of AMPK-inactivated and *Hk2*-deleted medulloblastomas, suggesting that both genes operate in a common pathway to maintain tumor stem cell populations.

## Discussion

Medulloblastomas are highly proliferative tumors that configure energy metabolism to support malignant growth through enhanced aerobic glycolysis [15, 16] and lipogenesis [17, 54]. AMPK is an intracellular energy sensor that coordinates anabolic and catabolic processes with nutrient availability and may crucially regulate tumor metabolism in ways that might be predicted either to support or to inhibit tumor growth. In SHH signaling, AMPK has been shown to interact directly with GLI1 to suppress SHH activity in medulloblastoma [55, 56]. In contrast to the anti-tumor effects of AMPK predicted by its interactions with GLI1, however, a prior study of SHH medulloblastoma showed that deletion of AMPK subunit *Prkaa2* slowed tumor progression in a primary mouse model [31]. Understanding the mechanisms through which AMPK inactivation reduces medulloblastoma growth may allow the design of targeted therapies that exploit the role of AMPK in SHH medulloblastoma and potentially in other cancers.

Our data show that AMPK inactivation slowed medulloblastoma growth and altered tumor cell heterogeneity by disproportionately impacting tumor stem cells. We inactivated AMPK by conditionally deleting both catalytic subunits, *Prkaa1* and *Prkaa2*, in mice carrying *Gfap-Cre* and *SmoM2* alleles, generating *G-Smo^Ampk^* mice with AMPK inactivation and SHH hyperactivation throughout the brain. In these mice, AMPK-inactivated medulloblastomas formed with 100% penetrance but progressed more slowly than medulloblastomas in controls with SHH hyperactivation but intact AMPK. Like control tumors, AMPK-inactivated tumors comprised medulloblastoma cells with a range of differentiation states. However, AMPK-inactivated medulloblastomas showed a shift in cell populations, with increased differentiated cells compared to controls and specifically reduced populations of OLIG2+ stem cells and malignant glia.

While AMPK inactivation affected different types of tumor cells in different ways, increasing differentiating populations and decreasing tumor stem cells, effects of gene expression were consistent across multiple cell types. In tumor cells across the differentiation spectrum, AMPK inactivation decreased the expression of multiple genes related to protein translation. This global down-regulation of translation-related genes suggested reduced mTORC1 activity, which was confirmed by the finding of reduced p4EBP1. Decrease in mTORC1 signaling in *G-Smo^Ampk^* tumors was unexpected, as acute AMPK activation inhibits mTOR [57, 58]. We speculate that *G-Smo^Ampk^* tumor cells reduce mTORC1 in a homeostatic response to AMPK inactivation.

*G-Smo^Ampk^* medulloblastomas also showed altered expression of glycolytic genes, with decreased *Hk2* and increased *Eno1*. By impeding the entry of glucose into the glycolysis through HK2 and enhancing the exit of glycolytic intermediates as phosphoenolpyruvate through ENO1, AMPK inactivation reduced the potential for diversion of glycolytic intermediates for other purposes.

We noted specific changes in the stromal populations in AMPK-inactivated tumors with important implications. Within the stromal populations subject to deletion of *Prkaa1* and *Prkaa2*, the less mature oligodendrocyte subset was specifically depleted in *G-Smo^Ampk^* medulloblastomas. These cells, like tumor stem cells, express OLIG2, and their decreased numbers suggests an interaction between AMPK and OLIG2 may crucially modulate OLIG2 function.

Within the stromal populations that were not subject to *Prkaa1* and *Prkaa2* codeletion, the endothelial cells were significantly increased in *G-Smo^Ampk^* medulloblastomas. As these cells were not within the GFAP-lineage, their increased population cannot be cell autonomous. Increased endothelial populations may indicated changes in tumor vasculature caused by AMPK inactivation in tumor cells, as we have previously seen in glycolysis deficient medulloblastomas [15]. This effect on endothelial cells may thus be another point of similarity between *Hk2*-deleted and AMPK-inactivated medulloblastomas.

By reducing stem cell self-renewal, glial trans-differentiation, HK2-dependent aerobic glycolysis and mTORC1 activation, AMPK inactivation altered multiple processes important for tumor progression. Each of these processes has been tested in isolation in previous studies. The maintenance of OLIG2+ stem cell pools has been shown to promote both medulloblastoma progression and recurrence after cytotoxic therapy [8]. HK2-dependent aerobic glycolysis is similarly required for tumor progression [15]. Paracrine signaling between malignant glia and myeloid cells within medulloblastomas also promotes tumor growth [11]. The activation of mTORC1 is similarly essential for tumor growth, and mTORC1 inhibition slows progression [59]. All of these processes are likely to be interrelated, and to contribute to the anti-tumor effect of AMPK inactivation. AMPK inactivation thus sets in motion a complex set of processes that result in the observed shift from multipotent stem cells to more differentiated tumor cells, with the net effect of slowing tumor growth.

To define causal relationships between processes that we found to be altered in AMPK-inactivated tumors, we compared the phenotypes of AMPK-inactivated and *Hk2*-deleted tumors. Both *Hk2* deletion and *Prkaa1/Prkaa2* co-deletion slow tumor growth and extend event-free survival. Importantly, *Hk2* deletion impaired stem cell maintenance and increased differentiation, reproducing the altered cellular heterogeneity of the *Prkaa1/Prkaa2* co-deleted tumors. These genetic data provide evidence for a linear pathway in which AMPK and HK2 both operate in the same direction to maintain undifferentiated, pluripotent populations within medulloblastomas.

The effect of AMPK inactivation on OLIG2+ stem cells identifies a vulnerability that may be exploited therapeutically for clinical benefit. Tumor stem cells are intrinsically resistant to cytotoxic therapy and drive recurrence. Ways to target these stem cells are urgently needed. Our data show that chronic blockade of AMPK activity disproportionately affects these stem cells, suggesting a way to target these otherwise resistant populations. Follow up studies are needed to test pharmacologic AMPK inhibition in combination with current, cytotoxic therapy. As genetic deletion studies demonstrate that AMPK inactivation can suppress tumor growth, and AMPK deletion in model organisms does not have clear deleterious effects [29], AMPK inhibition may emerge as an important new avenue for cancer therapy.

## Data Availability

The scRNA-seq data were deposited in the Gene Expression Omnibus database under the accession code GSE150579 and GSE GSE190297.

## Acknowledgements

We thank the UNC CGBID Histology Core, supported by P30 DK 034987 and the UNC Tissue Pathology Laboratory Core supported by NCI CA016086. T.D. was supported by NINDS (F31 NS120459). T.R.G. was supported by NINDS (R01NS088219, R01NS102627, R01NS106227) and by the UNC Department of Neurology Research Fund, and by a TTSA grant from the NCTRACS Institute, which is supported by the National Center for Advancing Translational Sciences (NCATS), National Institutes of Health, through Grant Award Number UL1TR002489.

## Author Contributions

T.D., D.S.M., B.D., A.T. and T.R.G. wrote the manuscript. T.D., D.S.M. and H.L. conducted the experiments and analyzed the data.

