## Supplementary Figure 1 for "Chronic AMPK inactivation slows SHH medulloblastoma progression by inhibiting mTORC1 signaling and depleting tumor stem cell populations"

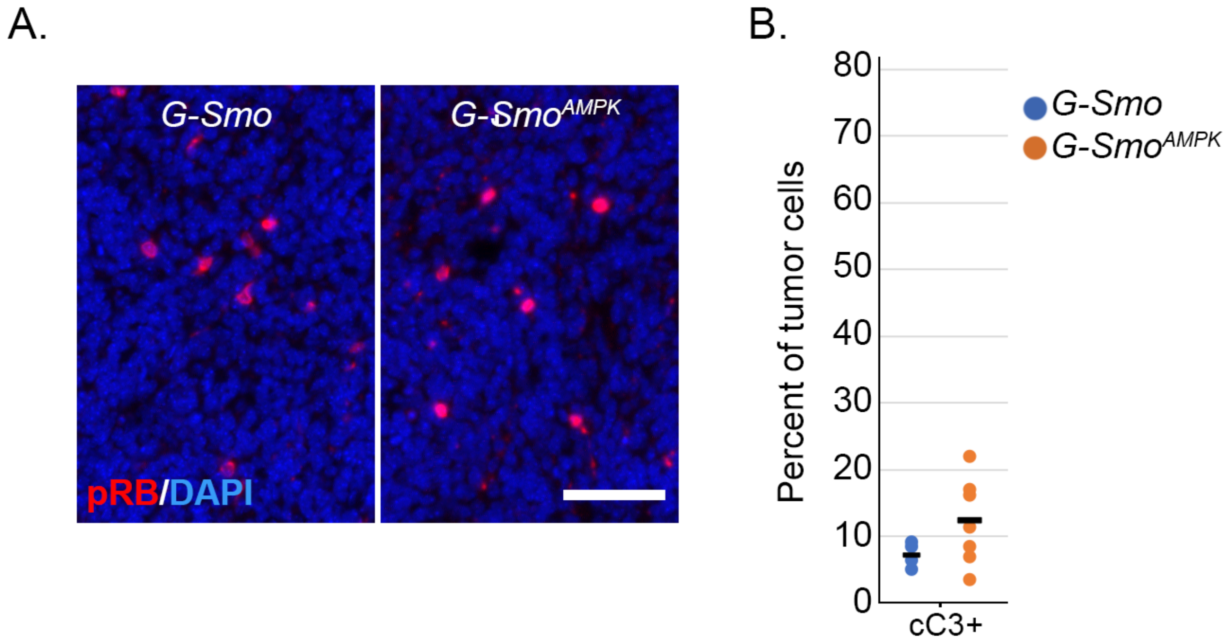


**Supplementary Figure 1. cC3 studies in *G-Smo*$^{AMPK}$ medulloblastomas and *G-Smo* controls.** (A) Representative IHC on sagittal sections of *G-Smo* and *G-Smo*$^{AMPK}$ medulloblastomas, showing cC3. (B) quantification of the fractions of cC3+ tumor cells in replicate tumors of each genotype.
